# Tracking the Canonical Computations of Perceptual Processing in a Quasi-Continuous Sensory Flow

**DOI:** 10.1101/232587

**Authors:** Jean-Rémi King, Valentin Wyart

## Abstract

The canonical computations involved in sensory processing, such as neural adaptation and prediction-error signals, have mainly derived from studies investigating the neural responses elicited by a single stimulus. Here, we test whether these computations can be tracked in a quasi-continuous flow of visual stimulation, by correlating scalp electroencephalography (EEG) recordings to simulations of neuronal populations. Fifteen subjects were presented with ~5,000 visual gratings presented in rapid sequences. Our results show that we can simultaneously decode, from the EEG sensors, up to 4 visual stimuli presented sequentially. Temporal generalization and source analyses reveal that the information contained in each stimulus is processed by a “visual pipeline”: a long cascade of transient processing stages, which can overall encode multiple stimuli at once. Importantly, our data suggest that the early feedforward activity but not the late feedback responses are marked by an adaptation phenomenon. Overall, our approach demonstrates how theoretically-derived computations, as isolated in single-stimulus paradigms, can be generalized to conditions of a continuous flow of sensory information.

## Introduction

The neuroscience of perception faces a challenge. On the one hand, our brain deals with a continuous and rapidly changing flow of sensory information. While on the other hand, the majority of studies decomposing the neural mechanisms of perceptual processing are based on the presentation of discrete stimuli. This single-stimulus paradigm has proved fruitful: a number of “canonical” computations such as predictive error signals, neural adaptation and normalization have been empirically established across a variety of sensory modalities and tasks (e.g. Carandini & Heeger 2012). However, it remains to be determined whether these computations account for, and generalize to, the processing of a continuous flow of sensory input. We make a step forward in addressing this question by tracking several theoretically-derived computations using EEG recordings of subjects presented with a rapid flow of visual stimulation.

## Methods

15 subjects were exposed to visual gratings in rapid sequences (Fig. 1), while their brain activity was recorded with 32-channel scalp electroencephalography (EEG). Data were low-pass-filtered at 16 Hz, epoched for each sequence of 8 items, and baseline corrected prior to the first stimulus onset. To quantify the extent to which i) the orientation of each grating (S) and ii) the differential of each grating (Δ_n_=S_n_ − S_n-1_) could be decoded from the EEG recordings at each time sample, we fit linear models as described in King et al. (2016), using the default parameters specified in MNE (Gramfort et al 2014). Temporal generalization analyses (King & Dehaene 2014) were based on the above linear models, and compared to stimulations of integrate-and-fire neuronal populations tuned to visual gratings. Simulations varied in terms of macroscopic architectures: i) random, ii) purely feedforward or iii) feedforward and feedback connections, as well as dynamical properties (with or without adaptation) using NEST& its standard parameters (Kunkel et al 2017).

**Figure 1.**
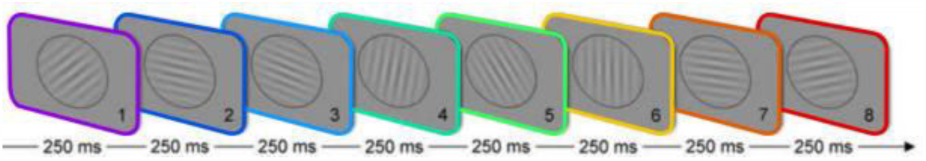
Subjects viewed rapidly changing gratings.

## Results

### Visual content and content change can be tracked with EEG

Our results show that subjects’ evoked response to each stimulus was correlated with both the orientation of the visual grating and the change of orientation between two subsequent gratings – which can be a proxy for prediction error signals (“Δ”, Fig. 2). Topographic and source analyses suggest that these two visual features are encoded in the visual and temporal cortices. Following the predictive coding framework, Δ elicited a much larger neural response than orientations;

**Figure 2.**
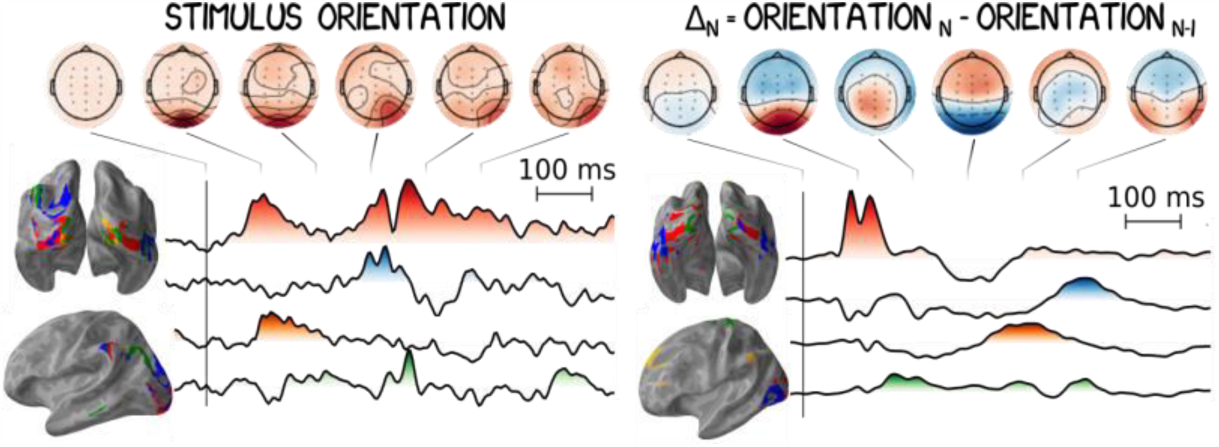
EEG codes for the orientation and Δ from ~90ms to 1,000 ms after grating onset.

### Sequentially presented stimuli can be simultaneously decoded from distinct processing stages

The orientation and the change of orientation in each sequential stimulus could be decoded for up to 1 second. Temporal generalization analyses (King & Dehaene 2014) confirmed that multiple stimuli can be simultaneously decoded at a unique time sample and further revealed strongly diagonal temporal generalization patterns (Fig. 3) – typical of perceptual processing. Sources estimation of the EEG patterns reveal the sequential recruitment of the early visual, lateral-occipital and prefrontal cortices. Together with neuronal simulations these EEG results suggests that visual processing is implemented by a neural pipeline: a hierarchical cascade of processing stages in which visual information is rapidly and continuously propagated. In this view, one snapshot of the entire cascade is sufficient to decode multiple stimuli, as each can been coded in distinct processing stages.

**Figure 3.**
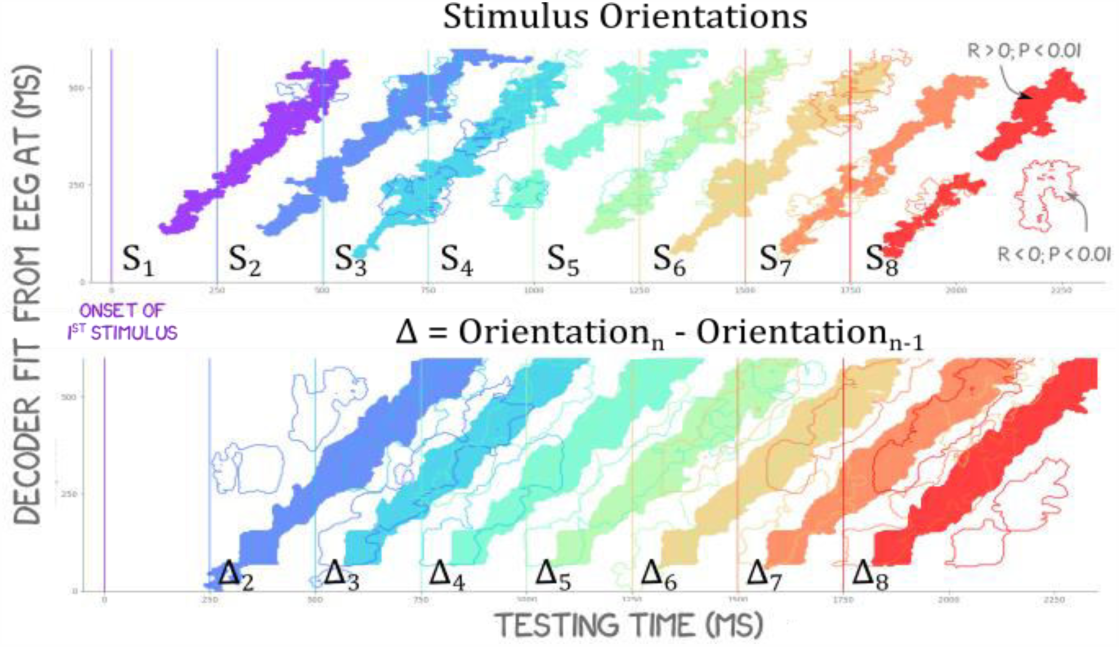
Fitting a decoder at each time point of the EEG activity and testing how it generalizes over time shows how each stimulus feature is propagated along a long cascade of processing stages.

### Early but not late processing stages are modulated by neuronal adaptation

We next quantified the extent to the processing stages coding for orientations, and orientation changes, interacted with one another. The results demonstrate a strong interaction between 90 and 350 ms: gratings that are more distinct from their preceding stimulus are encoded more strongly in early visual cortex as predicted by models of neuronal adaptation (Fig 4.). However, these adaptation effects decreased after 300 ms, suggesting that the late neural responses - marked by a topographical & source reversal (Fig 2.) typical of feedback - are less subject to adaptation. Overall, our results suggest that the feedforward but not the feedback of the visual pipeline is characterized by adaptation.

**Figure 4.**
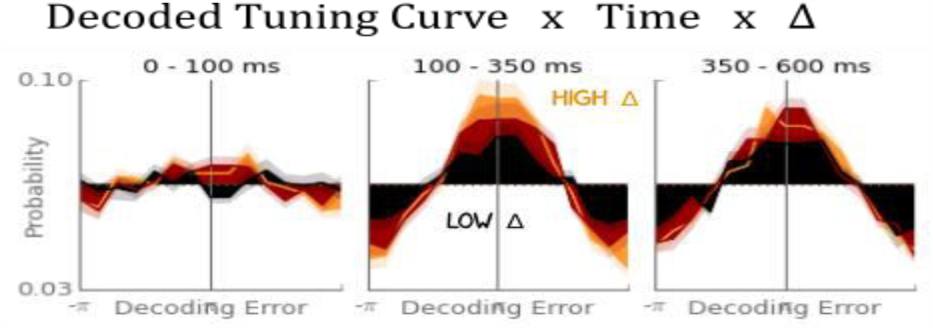
Adaptation, marked by enhanced codes with high Δ, is only present in early processing stages.

## Conclusion

We show that several theoretically-derived neuronal computations (encoding, prediction error & adaptation) can be tracked in a quasi-continuous flow of sensory stimulation. Additionally, our results i) reveal how sequential stimuli can be encoded simultaneously in distinct stages of a visual pipeline and ii) demonstrate that only the early stages are marked by neural adaptation. Overall, our approach opens the possibility to decipher the mechanisms of perceptual processing as they unfold under continuous sensory flow.

## Acknowledgments

European Union’s Horizon 2020 research &innovation program under the Marie Sklodowska-Curie Grant Agreement No. 660086, Bettencourt-Schueller and the Philippe Foundations (J-R.K).

